# Endothelial lipase facilitates low-density lipoprotein (LDL) uptake in LDL receptor deficiency by a heparan sulfate proteoglycan-dependent mechanism

**DOI:** 10.1101/2025.03.02.640965

**Authors:** Olivia White, Zahra Aligabi, Kendall H. Burks, Jingrong Tang, Nathan O. Stitziel, Ira J. Goldberg, Alan T. Remaley, Diego Lucero

## Abstract

Current lipid-lowering drugs are relatively ineffective in reducing low-density lipoprotein (LDL) cholesterol in patients with Familial Hypercholesterolemia (FH) due to a dysfunctional LDL receptor (LDLR). However, LDL cholesterol reductions have been achieved in FH patients using angiopoietin-like 3 (ANGPTL3) inhibitors, which act through an uncharacterized, LDLR-independent pathway that requires endothelial lipase (EL). Here, we aim to investigate EL’s direct role in LDLR-independent uptake of LDL in hepatocytes.

Control and LDLR-KO HepG2 cells were transfected with an empty plasmid or a plasmid encoding the human LIPG gene, and the cellular uptake of fluorescent human LDL was measured by FACS. To test the contribution of heparan sulfate proteoglycans (HSPG), LDL uptake was assessed with and without the pre-incubation with heparin or a heparinase cocktail. Finally, tetrahydrolipstatin was used to inhibit EL enzymatic activity in uptake studies.

Unsurprisingly, LDLR-KO HepG2 cells showed an 80% reduction in LDL uptake compared to controls (p<0.001). Remarkably, EL overexpression almost fully rescued LDL uptake in LDLR-KO cells (p<0.001), without effects in control cells. EL-mediated LDL uptake was completely blocked by heparinases and heparin in LDLR-KO cells, suggesting a crucial role of HSPG in EL-mediated LDL uptake. Notably, treatment with tetrahydrolipstatin reduced LDL uptake in EL-overexpressing LDLR-KO cells (p=0.0015). Our data reveals that EL facilitates the uptake of LDL in hepatocytes through an LDLR-independent, HSPG-dependent pathway that involves EL’s enzymatic activity. This pathway provides an additional mechanism to explain the reduction of LDL cholesterol induced by ANGPTL3 inhibitors and represents a potential druggable target to treat FH.

## INTRODUCTION

Familial hypercholesterolemia (FH) is characterized by elevated plasma levels of low-density lipoprotein (LDL) cholesterol (LDL-C), which causes accelerated atherosclerosis and leads to premature cardiovascular events (1). In most of the genetically characterized cases, FH is caused by autosomal dominant loss-of-function mutations in the LDL receptor (*LDLR*) gene and, less frequently, by mutations in other genes, such as apolipoprotein (APO) B, proprotein convertase subtilisin/kexin type 9 (*PCSK9*), and *APOE* (2). Regardless of the mutation involved, severe hypercholesterolemia in FH results from the deficient removal of LDL from circulation due to defects in the LDLR pathaway. In general, cardiovascular risk reduction is achieved in most patients by enhancing LDL clearance and lowering its plasma levels. The most commonly used LDL-lowering therapies, such as statins and PCSK9 inhibitors, lower plasma LDL by upregulating the LDLR pathway; therefore, their efficacy is limited in patients with low or null LDLR activity (3). Consequently, there is a need for new therapies that use alternative, LDLR-independent pathways to reduce plasma LDL-C in FH.

Inhibition of angiopoietin-like 3 (ANGPTL3), a member of the ANGPTL family, has recently been shown to reduce LDL-C in patients with a homozygous loss of the LDLR (4, 5). ANGPTL3 is expressed in the liver and secreted to the circulation, where it inhibits the activity of lipoprotein lipase (LPL) and endothelial lipase (EL), key enzymes in the metabolism of triglyceride-rich lipoproteins and high-density lipoproteins (HDL), respectively (6). In an effort to develop new therapies to reduce plasma triglycerides, ANGPTL3 has been targeted with monoclonal antibodies (5, 7), antisense oligonucleotides (8), and silencing RNA (9). Unexpectedly, ANGPTL3 inhibition led to a marked reduction in LDL-C in homozygous FH patients with null LDLR activity (5, 10). Further studies in mice lacking LDLR showed that the ANGPTL3 inhibitor-induced LDL reduction requires the presence of EL (11, 12). Nevertheless, the detailed mechanisms behind this LDLR-independent pathway for LDL removal remain obscure.

Endothelial lipase is a heparin-released lipase that is anchored on heparan sulfate proteoglycans (HSPGs) (13) and plays a key role in lipoprotein metabolism. For instance, EL mediates HDL catabolism either by phospholipolysis or by facilitating HDL-holoparticle uptake in hepatocytes (14, 15). Moreover, in murine models, EL deficiency causes accumulation of VLDL particles (16), whereas hepatic EL overexpression enhances the clearance of plasma LDL (17). More recently, a study in *Ldlr*-knockout mice demonstrated that EL is essential for the LDL-lowering effect of ANGPTL3 inhibitors in the absence of LDLR (12). The authors hypothesized that EL activation by ANGPTL3 inhibition promotes VLDL remodeling and clearance of VLDL remnants, preventing LDL formation (12). Nevertheless, evidence suggests that there are additional mechanisms at play in the ANGPTL3 inhibitor-mediated LDL reduction. In fact, human kinetic studies in FH patients treated with ANGPTL3 inhibitors revealed an increase in the plasma clearance of LDL particles (18). Based on these findings, we hypothesize that EL activation by ANGPTL3 inhibition may also play a direct role in LDL catabolism independently of the LDLR.

Here, we tested whether EL directly modulates the cellular uptake of LDL in hepatocytes by overexpressing EL in LDLR-expressing and LDLR-deficient hepatoma cells. This model enabled us to focus on EL’s actions on cellular LDL uptake independently of any pleiotropic effects from ANGPTL3 inhibition or deletion (19). We measured cellular LDL uptake, evaluated the role of EL enzymatic activity, and tested the contribution of HSPGs. Although EL does not impact LDLR-mediated LDL uptake significantly, we found that EL overexpression almost fully rescued cellular LDL uptake in LDLR-deficient hepatocytes. This process relies on EL’s enzymatic activity and involves EL binding to cell surface HSPGs. Our data highlights the existence of alternative pathways for hepatic LDL uptake independently of the LDLR and provides an additional mechanism that could explain the reduction of plasma LDL observed in FH patients treated with ANGPTL3 inhibitors.

## METHODS

### 1.- Data availability

The data from experiments leading to the most relevant findings is available on the NHLBI FigShare account: https://figshare.com/s/c372d7fc44d1ff124c75

### 2.- Reagents

HepG2 cell line (Cat. # T3256) were purchased from ABM Goods, Inc. (Canada). Alexa Fluor^TM^ 568 carboxylic acid, succinyl ester (Cat. # A20184), and DiI–human LDL conjugate (Cat. # L3482) were purchased from Molecular Probes, (Eugene, OR). Tetrahydrolipstatin (THL) (Cat. # O4139-25MG) was purchased from Sigma-Aldrich (St. Louis, MO). High-Capacity RNA-to-cDNA™ Kit (Cat. # 4388950), TaqMan Master Mix (Cat. # 4304437), and TaqMan Gene Expression Assays (LIPG [Hs00195812_m1] and 18S [Hs03003631_g1]) were obtained from Applied Biosystems (Waltham, MA). Lipofectamine 2000 (Cat. # 11668019) was purchased from Invitrogen^TM^ (Waltham, MA). The empty plasmid (Cat. # PS100001) and the plasmid containing human LIPG (Cat. # RC209248) were purchased from OriGene Technologies Inc. (Rockville, MD). Heparinases, Heparinase I (Part # 50-008), Heparinase II (Part # 50-011), and Heparinase III (Part # 50-012), were obtained from IBEX (Montreal, Canada). Rabbit anti-human LIPG primary antibodies (Cat. # ab24447) and HRP-conjugated Goat anti-Rabbit IgG secondary antibody (Cat. # ab6721) were purchased from Abcam (Cambridge, United Kingdom). Goat anti-human GAPDH polyclonal antibody (V-18) conjugated to HRP (Cat. # sc-20357) was purchased from Santa Cruz Biotechnology, Inc. (Dallas, TX). The EnzChek™ Phospholipase A1 Assay Kit (Cat. # E10219) was purchased from Invitrogen ^TM^ (Waltham, MA).

### 3.- Cell culture

A Cas9 stably expressing HepG2 cells (ABM Goods, Canada; Cat# T3256) were cultured in Dulbecco’s Modified Eagle Medium (DMEM) supplemented with 100 IU/mL of penicillin G, 100 µg/mL streptomycin and 10% (vol/vol) fetal bovine serum (FBS). Cells were incubated in a humidified incubator with 5% CO_2_/95% air and split every 2 or 3 days.

### 4.- Generation of LDLR knockout HepG2 Cells

LDLR-deficient HepG2 cells were generated using CRISPR/Cas9 as previously described (20). Briefly, stably Cas9-expressing HepG2 cells were transduced with lentivirus containing sgRNA targeting human LDLR (ABM Goods, Inc. Canada, Cat# 264181110204). Transduced cells were sorted into 96-well plates (one cell per well) in an BD FACSAria™ III Cell Sorter. Clones were expanded, and the loss of LDLR was tested at mRNA and protein levels by RT-qPCR and Western Blot, respectively. In addition, phenotype was confirmed by fluorescent LDL uptake, showing ∼80% reduction in LDL uptake.

### 5.- Overexpression of endothelial lipase

Control and *LDLR*-KO HepG2 cells were transfected with an empty plasmid or a plasmid containing human *LIPG* gene to enhance basal expression of EL. Cells were plated in 24-well plates in DMEM supplemented with 100 IU/ml of penicillin G, 100 µg/mL streptomycin and 10% (vol/vol) FBS, at a density of 40,000 cells/well. After overnight incubation for cell adherence, cells were then transfected with an empty plasmid or a plasmid containing the Myc-DDK-tagged human LIPG by lipofection using Lipofectamine 2000 following the manufacturer’s instructions (Invitrogen, Waltham, MA). After 72 hours, cells were processed for total RNA or protein extraction and cellular LDL uptake as described below.

### 6.- RT-qPCR

Total RNA from control and *LDLR*-KO HepG2 cells, transfected either with an empty plasmid or a LIPG-plasmid, was extracted using a Qiagen RNeasy® Mini Kit (Qiagen, Hilden, Germany), according to the manufacturer’s instructions.

A total of 1 µg of RNA was converted into cDNA using High-Capacity RNA-to-cDNA™ Kit (Applied Biosystems, Waltham, MA), following the manufacturer’s instructions. RT-qPCR was carried out in 384-well plates using TaqMan Gene Expression Assays and TaqMan Master Mix (Applied Biosystems, Waltham, MA), according to the manufacturer’s instructions.

### 7.- Transcriptomic analysis

The expression of *LIPG*, the gene encoding EL, in liver tissue-derived cells was determined using The Tabula Sapiens (21) and Human Protein Atlas (22) databases. UMAP plots of *LIPG* expression (EL) in liver cells sorted by ontology class from the Tabula Sapiens database were illustrated through CZ CellxGene Discover (23).

### 8.- Immunoblotting

Control and *LDLR*-KO HepG2 cells were plated in 6-well plates, at a density of 250,000 cells per well, and incubated for 24 h for cell adherence. Next, cells were transfected by lipofection with an empty plasmid or a plasmid containing the human *LIPG* gene. 72 h post-transfection, cells were washed and incubated with vehicle (PBS) or 15 U/mL of heparin for 6 h at 37°C, in Opti-MEM^TM^ Serum-free media (Gibco, Waltham, MA). Next, supernatant was collected, added Halt™ protease and Halt™ phosphatase inhibitors (Thermo Fisher Scientific, Waltham, MA), and centrifuged to eliminate cell debris. Cells were washed with ice-cold PBS and lysed with RIPA buffer in the presence of Halt™ protease and Halt™ phosphatase inhibitors (Thermo Fisher Scientific, Waltham, MA). Western blots were performed as previously described (24) using rabbit anti-human LIPG (1:500 and 1:2000) (Abcam, Cambridge, UK) and goat anti-rabbit antibody conjugated to HRP (1:10,000) (Abcam, Cambridge, UK) as secondary antibody. In cell lysate blots, HRP-conjugated anti-GAPDH (1:2,000) (Santa Cruz Biotechnology, Dallas, TX) was used as loading control.

### 9.- Preparation and fluorescent labeling of lipoproteins

Human LDL (*d*: 1.019 – 1.064 g/mL) was isolated from plasma from healthy donors by preparative sequential KBr density ultracentrifugation (330,000 x g) followed by extensive dialysis at 4° against PBS to remove KBr. Total protein content in LDL was determined using Pierce™ BCA Protein Assay Kits (ThermoFisher, Waltham, MA). Pure LDL (6 mg of protein) was then incubated with Alexa Fluor^TM^ 568 carboxylic acid, succinyl ester (Molecular Probes, Eugene, OR) for 1 hour at room temperature in the dark. Free, unbound dye was separated from labeled LDL by preparative Fast Protein Liquid Chromatography (FPLC) using a HiTrap Desalting column (GE Healthcare, Chicago, IL). Labeled LDL was collected in the void volume and the LDL fraction’s protein concentration was reassessed using the above-mentioned method.

### 10.- Cellular LDL uptake assays

Cellular LDL uptake was assessed as previously described with slight modifications (20). Briefly, 72 hours post transfection, control and *LDLR*-KO HepG2 cells were incubated with 20 µg/mL of fluorescently labeled LDL (Alexa Fluor^TM^ 568-LDL) in serum-free DMEM containing 0.1% bovine serum albumin (Sigma-Aldrich, St. Louis, MO) for 90 minutes at 37°C in the same 24-well plates. Cells were then washed with warm PBS and dissociated using 0.25% trypsin (Gibco, Waltham, MA). Cells were then resuspended in ice-cold PBS containing 0.5% BSA and 2.5 µM EDTA and transferred to 96-well plates for flow cytometry analysis. Mean intensity of cell-associated Alexa Fluor^TM^ 568 fluorescence, proportional to cellular LDL uptake, was measured by fluorescence activated cell sorter (FACS) flow cytometry in a BD LSRFortessa (BD Biosciences, Franklin Lakes, NJ). The number of counted events was at least 10,000 for each measurement.

In the competition studies, cells were incubated with 20 µg/mL of fluorescently labeled LDL (Alexa Fluor^TM^ 568-LDL) in the presence of an excess (20X) of unlabeled LDL (400 µg/mL). Cellular LDL uptake was measured as indicated above.

#### 10.a Heparin treatment

A set of control and *LDLR*-KO HepG2 cells, transfected with either the empty plasmid or *LIPG*-plasmid, were pre-treated with 5 U/mL of heparin for 30 minutes at 37°C immediately before the cellular LDL uptake experiment. Cells were washed twice with warm PBS and processed to assess the cellular LDL uptake, as indicated above. In a parallel experiment, cells did not receive heparin pre-treatment, but the cellular LDL uptake assay was performed in the presence of 5 U/mL of heparin for 90 minutes at 37°C.

#### 10.b Heparinase treatment

Immediately before the cellular LDL uptake assay, control and *LDLR*-KO HepG2 cells, either transfected with the empty plasmid or *LIPG*-plasmid, were treated with vehicle (PBS) or a heparinase mix (Heparinase I 2.5 mU/mL; Heparinase II 2.5 mU/mL; Heparinase III 5 mU/mL; IBEX, Montreal, Canada) for 30 minutes at 37°C, in 0.5% BSA DMEM (25). Next, cells were washed twice with warm PBS and processed to assess the cellular LDL uptake, as indicated above.

#### 10.c Tetrahydrolipstatin treatment

Before the cellular uptake assays, control and *LDLR*-KO HepG2 cells, either overexpressing *LIPG* or not, were incubated with either 25 µg/mL of tetrahydrolipstatin (THL) (Sigma-Aldrich, St. Louis, MO) or the equivalent amount of vehicle (DMSO) for 3 hours in complete media at 37°C to inhibit EL activity. Next, the cellular LDL uptake assay was performed as indicated above in the presence of vehicle or THL (25 µg/mL).

### 11.- Time-course LDL uptake

Control and *LDLR*-KO HepG2 cells were seeded in 24-well plates at a 40,000 cell/well density, transfected as indicated above to overexpress *LIPG*, and incubated in complete growth media for 72 h. After that, cells were chilled to 4°C, washed with ice-cold, and incubated for 1 h at 4°C in 0.5% BSA in DMEM containing 15 µg/mL of pHrodo™ Red-LDL Conjugate (Invitrogen, Waltham, MA), a pH-sensitive dye that strongly fluoresces at endosomal pH. Next, cells were washed with ice-cold PBS to eliminate unbound LDL and then incubated at 37°C for 10, 20, 30, and 45 minutes to allow LDL internalization. Cells were then washed with ice-cold PBS and incubated with ice-cold 15 U/mL heparin in PBS to remove the remaining LDL on the cell surface. Finally, cells were dissociated with 0.25% trypsin, resuspended in FACS buffer, and transferred to a 96-well plate for analysis. Mean intensity of cell-associated pHrodo red fluorescence was measured by FACS flow cytometry in a BD LSRFortessa (BD Biosciences, Franklin Lakes, NJ). The number of counted events was at least 10,000 for each measurement.

### 12.- Phospholipase activity assay

Control and *LDLR*-KO HepG2 cells were seeded in 24-well plates at a 40,000 cell/well density, transfected as indicated above with an empty plasmid or a plasmid containing the human *LIPG* gene, and incubated in complete growth media for 72 h. Cells were pre-incubated with 25 µg/mL THL or vehicle in complete growth media for 3 h at 37°C. After that, cells were incubated in 5 U/mL heparin in Opti-MEM^TM^ Serum-free media (Gibco, Waltham, MA) for 90 minutes at 37°C to detach the EL from HSPGs. THL (25 µg/mL) and vehicle were kept in the media throughout the experiment. Then, conditioned media was collected on ice, centrifuged to remove cells and cell debris, and immediately processed for phospholipase activity. Phospholipase A1 activity was measured using the EnzChek™ Phospholipase A1 Assay Kit (Invitrogen, Waltham, MA), following the manufacturer’s instructions. Data is presented as U/µL of conditioned media.

### 13.- LDL binding to cell surface

Control and *LDLR*-KO HepG2 cells, overexpressing LIPG or not, were chilled down to 4°C for 1 h, and then incubated with 20 µg/mL of DiI-LDL complex in 0.5% BSA DMEM for 1 h at 4°C. Then, cells were washed with ice-cold PBS containing calcium and magnesium and dissociated by pipetting up and down in ice-cold PBS, containing calcium and magnesium. Mean intensity of cell-associated DiI fluorescence was measured by FACS flow cytometry in a BD LSRFortessa (BD Biosciences, Franklin Lakes, NJ). The number of counted events was at least 10,000 for each measurement.

### 14.- Statistical analysis

Data are presented as mean ± SD. Data distribution was verified with the Shapiro-Wilk test. Unless otherwise indicated, differences between groups were tested by Student’s t-test and One-way ANOVA, followed by Dunnett’s test, depending on each experiment design, and two-sided P < 0.05 were considered statistically significant. Data was analyzed in Prism 10. Simple linear regression was applied to the time-course LDL uptake studies. Data is presented as Slope [95% CI]. Model pair comparisons were tested in Prism 10.

## RESULTS

Despite its name, early studies demonstrated that EL is also expressed in liver tissue (26); however, the cellular distribution of hepatic EL expression is unknown. Our analysis of publicly available single-cell transcriptomic databases from human liver revealed that in addition to endothelial cells in the liver, hepatocytes themselves express significant amounts of *LIPG* mRNA (Figure 1A).

**Figure 1:**
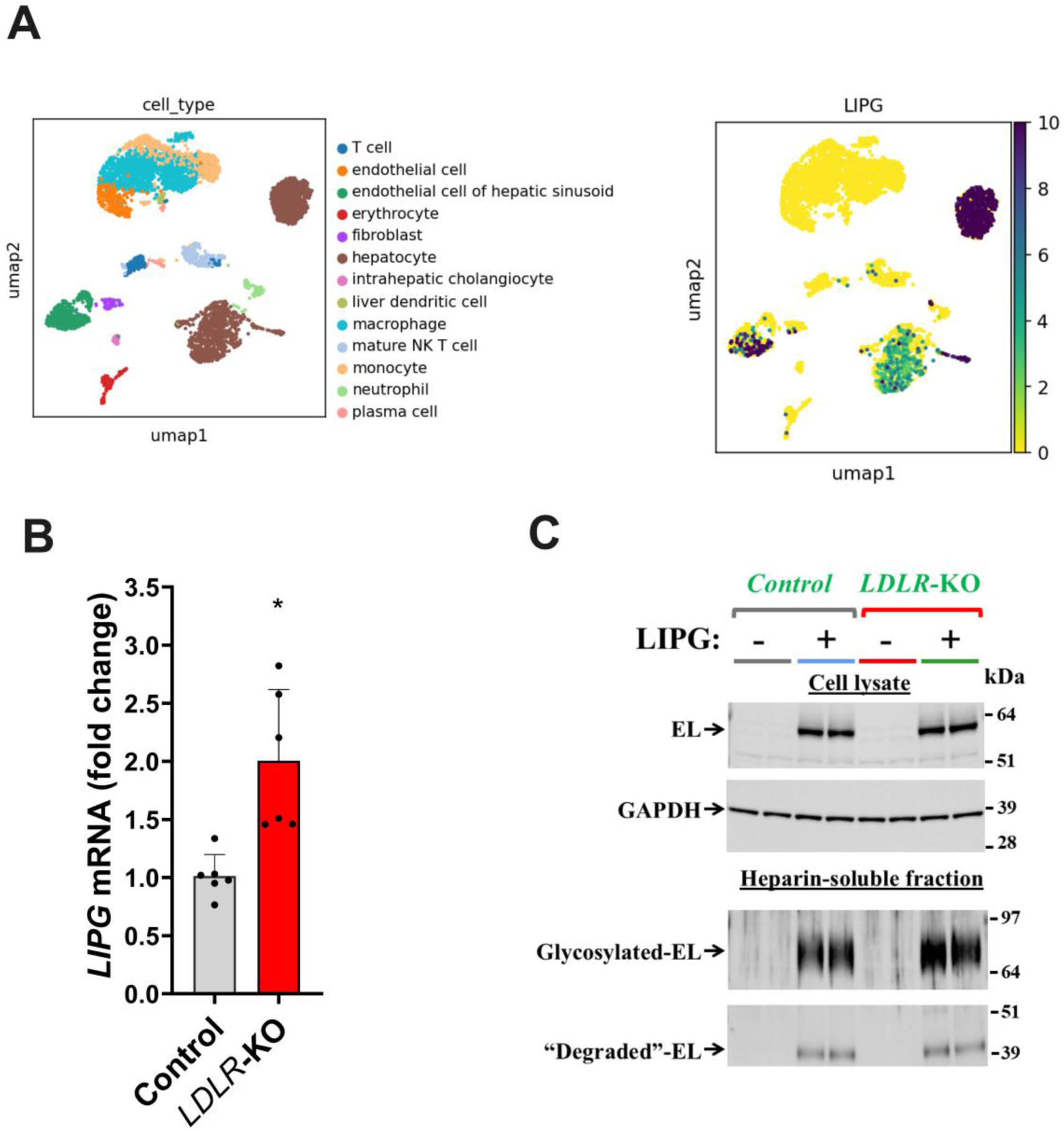
Hepatocytes show high LIPG expression levels, and the expression is upregulated by low-density lipoprotein receptor (LDLR) deficiency. Overexpressing LIPG in HepG2 cells resulted in heparan sulphate proteoglycan-associated expression of endothelial lipase (EL). A. Relative expression of LIPG in human hepatocytes and endothelial cells. B. Relative LIPG expression in control and LDLR-knockout (KO) HepG2 cells. Cells were plated and incubated with 20 µg/mL of LDL for 48 h, total RNA was extracted, and the levels of LIPG mRNA was measured by RT-qPCR. Data is presented as mean ± SD. Student’s T-Test. *p=0.0035 vs. Control. C: Endothelial lipase protein expression in EL-transfected control and LDLR-KO HepG2 cells. Cells were transfected with an empty plasmid or a plasmid containing human LIPG. After 72 h, cells incubated for 6 h with media containing 15 U/mL of heparin. Then, conditioned media (heparin-soluble fraction) and cell lysates were collected, and EL expression was assessed by immunoblotting.

To study EL’s role in modulating cellular LDL uptake in liver cells, we used control and *LDLR*-KO hepatoma cells (HepG2) previously developed in our laboratory (20). Although HepG2 cells presented relatively low *LIPG* mRNA expression levels, the loss of LDLR in HepG2 cells resulted in a 2-fold increase in *LIPG* mRNA levels (p=0.0035 vs. control cells; Figure 1B). Nevertheless, given their low basal EL expression, HepG2 cells represent an advantageous model to study the effects of EL overexpression with minimal interference of endogenous EL expression. We transfected control and *LDLR*-KO cells with an empty plasmid (baseline *LIPG* expression) or a plasmid harboring the human *LIPG* gene. Our protein expression analysis confirmed the presence of EL in the cell lysate and the heparin-soluble fraction (Figure 1C), indicating that EL is secreted and resides anchored to HSPGs on the cell surface. The molecular weight shift, from 55 to 68 kDa, observed between intracellular and heparin-released EL suggests that HSPG-bound EL is heavily glycosylated.

To investigate EL’s role in cellular LDL internalization, we assessed the capacity of the cells to internalize fresh, fluorescently labeled human LDL. In line with the loss of the LDLR, the cellular uptake of LDL was greatly decreased in *LDLR*-KO cells (p<0.0001; Figure 2A), with the residual uptake (∼20%) being mediated by LDLR-independent pathways. Remarkably, EL overexpression had opposing effects on the cellular LDL uptake in control and *LDLR*-KO HepG2 cells (Figure 2B and 2C). While control cells exhibited a slight reduction in cellular LDL uptake (−16%; p<0.0001; Figure 2B), LDLR-deficient cells showed an increase in cellular LDL uptake of almost 2-fold in response to EL overexpression (p<0.0001; Figure 2C). These results indicate that in the absence of a functional LDLR, EL promotes LDL uptake in liver cells through an LDLR-independent mechanism.

**Figure 2.**
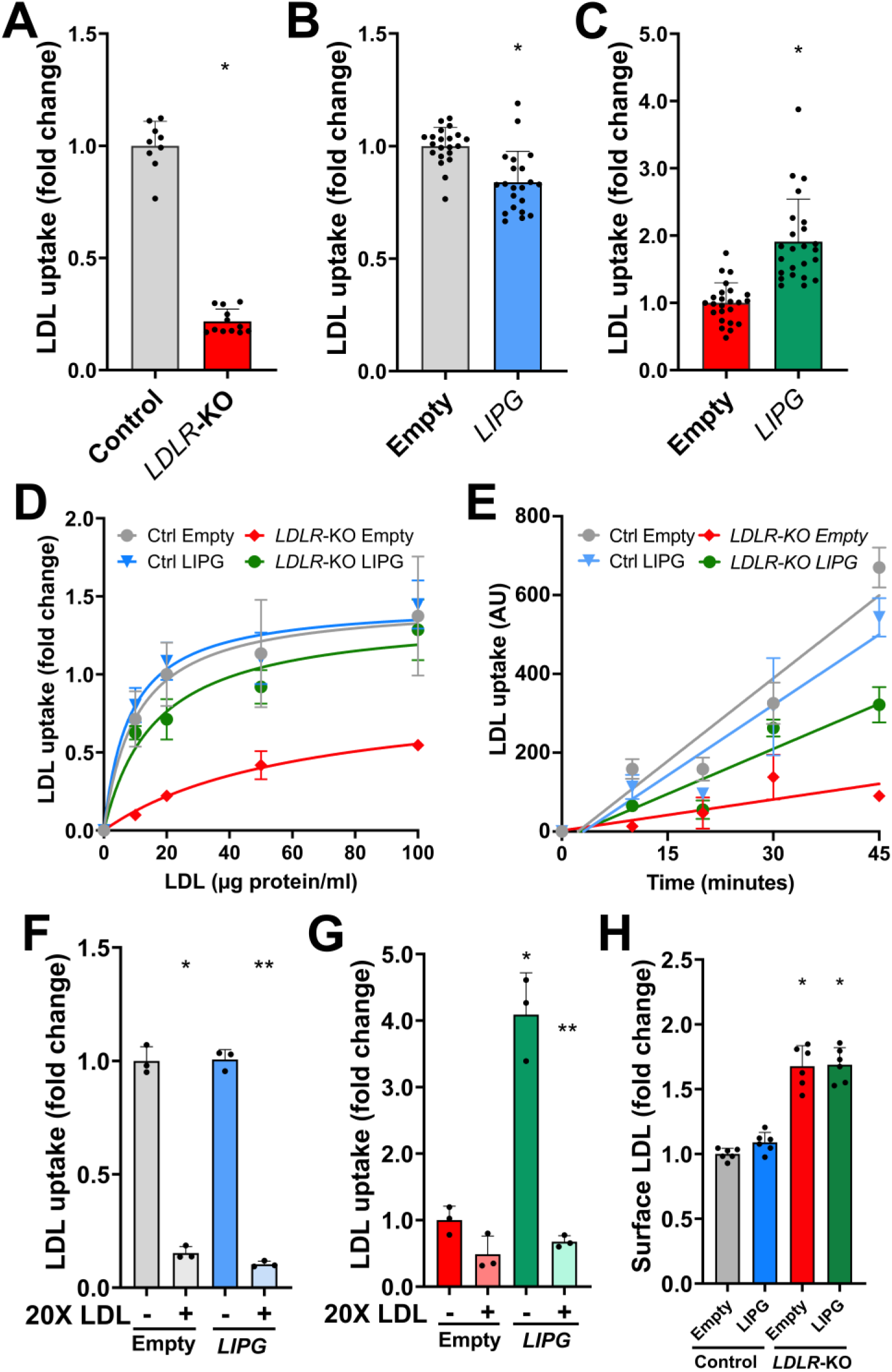
Endothelial lipase (EL) overexpression increases the cellular uptake of LDL by enhancing the LDL internalization rate only in LDLR-deficient hepatoma cells through a saturable pathway without affecting the LDL binding to cell surface. A: Cellular LDL uptake in control and LDLR-KO HepG2 cells. Control and LDLR-KO HepG2 cells were incubated with fluorescently labeled LDL (20 µg/ml) for 90 minutes. Cell-associated fluorescence was measured by FACS. Data is presented as mean ± SD. Student’s T-Test. *p<0.001. B and C: Cellular LDL uptake in control (B) and *LDLR*-KO (C) HepG2 cells, with or without EL overexpression (LIPG). Cells were transfected with either an empty plasmid or a plasmid containing the human LIPG gene (EL). After 72 h, cells were tested for cellular LDL uptake as previously indicated. Data is presented as mean ± SD. Student’s T-Test. *p<0.001. D: Cellular LDL uptake vs LDL concentration in control and *LDLR*-KO HepG2 cells with and without *LIPG* overexpression. Cells were incubated with the indicated concentrations of fluorescent LDL for 90 minutes. Uptake was analyzed by FACS. E: Time-course cellular LDL uptake in control and LDLR-KO HepG2. Cells were transfected with either an empty plasmid or a plasmid containing the human LIPG gene (EL). After 72 h, cells were pre-chilled and incubated with 15 ug/ml pHrodo-LDL (Invitrogen, Waltham, MA) for 1 h at 4°C. Cells were then thoroughly washed and incubated at 37°C for 10, 20, 30, or 45 minutes. LDL uptake, proportional to pHrodo fluorescence, was quantified by FACS. Lines represent the linear regression analysis. F and G: LDL uptake competition assay of fluorescently labeled LDL in the presence of excess (20X) unlabeled LDL in control (F) and LDLR-KO (G) HepG2 cells, with and without EL overexpression (LIPG). Cells were incubated with 20 µg/mL of fluorescently labeled LDL in the presence of 400 µg/mL unlabeled LDL for 90 minutes. Uptake was measured by FACS. One-way ANOVA. *p<0.0001 vs cells transfected with empty plasmid without excess of LDL. ** p<0.0001 vs LIPG-overexpressing cells without excess of LDL. H: Quantification of LDL bound to cell surface. Pre-chilled cells were incubated with 20 ug/ml of LDL for 1 h at 4°C. Then, cells were thoroughly washed with ice-cold PBS and cell associated fluorescence was measured by FACS. One-way ANOVA. *p<0.0001 vs control cells transfected with empty plasmid.

To further characterize the effects of EL overexpression on LDL uptake, cells were exposed to either increasing concentrations of LDL at a fixed time point (90 minutes) or increasing incubation times at a fixed LDL concentration. Figure 2D shows the cellular LDL uptake in response to various LDL concentrations normalized to that in control cells transfected with empty plasmid at 20 µg/ml LDL. Notably, EL-overexpression in *LDLR*-KO almost fully rescued the cellular LDL uptake, suggesting a more dynamic LDL uptake in the presence of EL. In fact, our time-course study revealed a steeper LDL internalization curve in the *LDLR*-KO HepG2 cells that overexpressed EL compared to those transfected with an empty plasmid (Slope [95% CI]: 7.6 [5.5 – 9.8] vs 2.6 [0.6 – 4.7]; F=13.2, p=0.0012) (Figure 2E). In contrast, EL overexpression did not change the rate of LDL internalization in control cells (Slope [95% CI]: 14.0 [10.8 – 17.2] (Empty) vs 11.9 [8.0 – 15.9] (*LIPG*); F=0.8, p=0.37) (Figure 2E). As LDL uptake in EL-overexpressing *LDLR*-KO cells saturates at relatively low LDL concentrations (Figure 2D), we implemented a competition assay to characterize the EL-mediated LDL internalization process by assessing LDL uptake in the presence of 20X unlabeled LDL. As expected, excess of unlabeled LDL strongly competed with labeled LDL for the LDLR-dependent uptake in control cells regardless of EL expression levels (p<0.0001, Figure 2F). Remarkably, while little effect was observed in *LDLR*-KO cells without EL overexpression (Figure 2G), excess of unlabeled LDL strongly inhibited the uptake of fluorescent LDL in the *LDLR*-KO cells that overexpressed EL (Figure 2G), similar to LDLR-mediated uptake. These findings suggest that, in LDLR deficiency, EL induces LDL internalization through a pathway with relatively limited capacity. Next, to interrogate whether the enhanced dynamics of LDL internalization was due to more avid binding of LDL to the cell surface, we incubated cells with fluorescent LDL in the cold.

*LDLR*-KO cells showed enhanced LDL binding to the cell surface than control cells (p<0.05) (Figure 2F), which may be due to compensatory mechanisms activated in response to LDLR deficiency. However, LDL binding was not affected by EL overexpression (Figure 2F). These findings suggest that in *LDLR*-KO cells, the presence of EL is required to enhance LDL internalization, although LDL may not bind directly to EL.

Endothelial lipase is known to be attached to the HSPGs on the cell surface (Figure 1E). This led us to investigate the role of HSPGs in the EL-mediated cellular LDL uptake. We tested whether heparin (5 U/ml) affected the uptake of LDL. Heparin treatment of control cells led to a slight increase in LDL uptake regardless of EL overexpression (Figure 3A and C). In contrast, both co-incubation and pre-incubation with heparin strongly inhibited the uptake of LDL in *LDLR*-KO cells (p<0.03; Figures 3B and 3D). Remarkably, heparin’s effect was similar in cells with and without EL overexpression. To confirm that these results were due to changes in cell surface HSPGs, we pre-incubated the cells with heparinases, which eliminate the side chains of HSPGs. Cellular LDL uptake was unaffected by heparinase pre-incubation in control cells (p>0.98; Figure 3C). Conversely, like heparin treatment, heparinases markedly decreased cellular LDL uptake in *LDLR*-KO cells, regardless of the overexpression of EL (p<0.0013; Figure 3F). Our data suggest that when EL is overexpressed in LDLR deficient cells, LDL uptake is mediated by the HSPG-bound EL fraction. Furthermore, HSPGs seem to play a fundamental role in the *LDLR*-KO cells’ residual LDL uptake.

**Figure 3.**
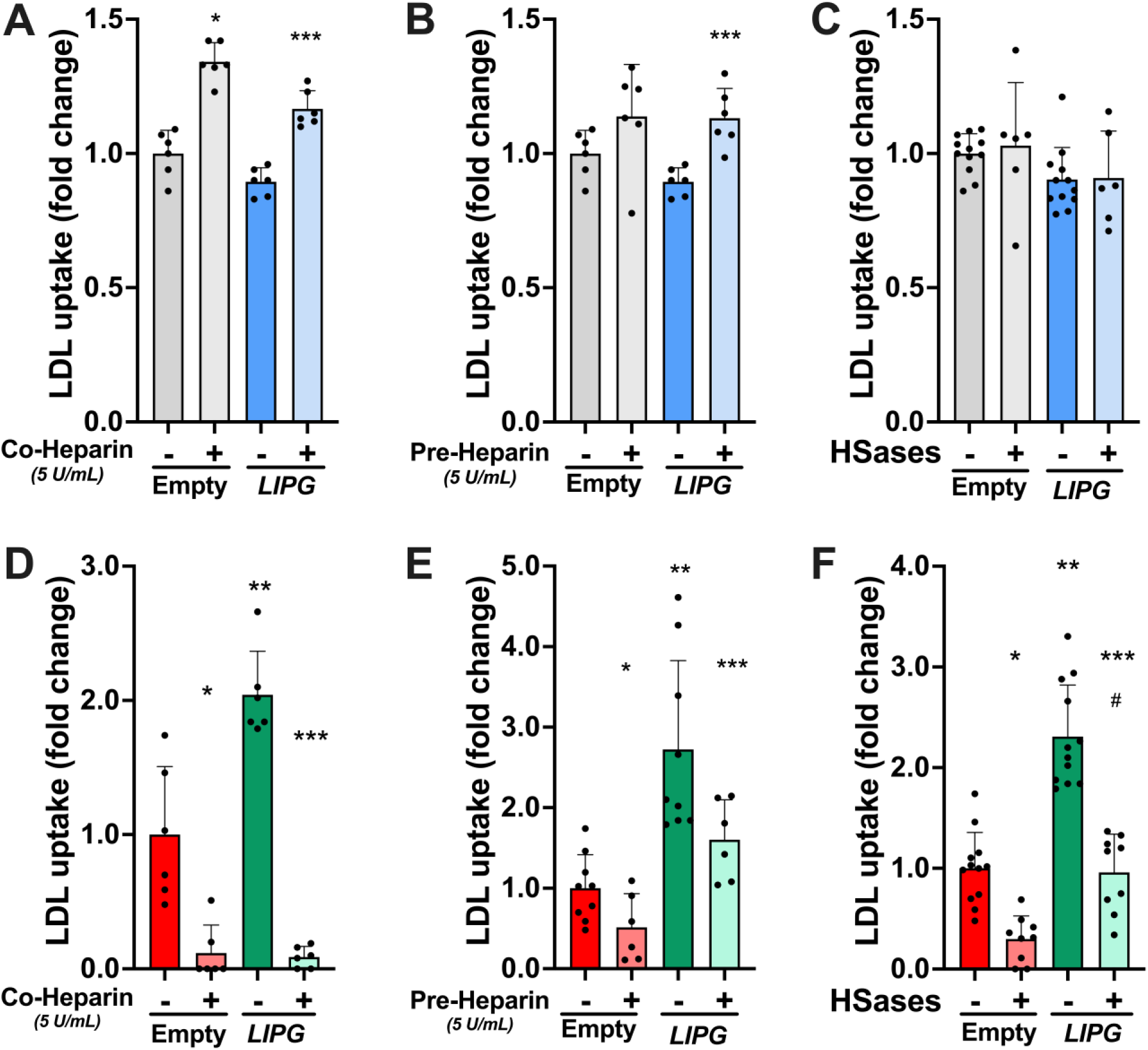
The disruption of the heparan sulfate proteoglycans alters the endothelial lipase (EL)-mediated cellular LDL uptake in LDLR-deficient hepatoma cells. Cellular LDL uptake was assessed in control (A, B, and C) and LDLR-KO (D, E, and F) HepG2 cells, with or without EL overexpression (LIPG). Cells were incubated with 20 ug/mL fluorescent LDL for 90 minutes either in the presence of vehicle (PBS) or 5 U/mL heparin, or after the incubation for 30 minutes at 37°C with vehicle (PBS) or 5U/mL heparin or heparinase cocktail (Heparinase I and II at 2.5 mU/ml, and Heparinase III at 5 mU/ml; IBEX, Montreal, Canada). After the pre-incubation, no heparin or heparinases were present during the uptake study. Cellular LDL uptake was quantified by FACS. A and D: Cellular LDL uptake in the presence heparin. B and E: Cellular LDL uptake after heparin pre-incubation. C and F: Cellular LDL uptake after heparinase treatment. Data is presented as mean ± SD. One-way ANOVA. *p<0.030 vs. cells transfected with empty plasmid plus vehicle. **p<0.002 vs. cells transfected with empty plasmid plus vehicle. ***p<0.025 vs. LIPG-overexpressing cells plus vehicle. #p<0.0001 vs. LIPG-overexpressing cells plus Heparinases. Hases: Heparinases.

To elucidate whether EL enzymatic activity is required for its influence on cellular LDL uptake, we tested whether a panlipase inhibitor, THL, would reduce LDL uptake in *LDLR*-KO cells. As expected, based on our previously shown data, THL did not change the uptake in control cells (Figure 5A). In contrast, the inhibition of enzymatic activity with THL reduced the uptake in the *LDLR*-KO cells that overexpressed EL (p=0.0015), while THL treatment did not affect the uptake in the *LDLR*-KO cells transfected with the empty plasmid (p=0.98). Importantly, THL effectively inhibited EL activity as confirmed by measuring A1 phospholipase activity in the post-heparin supernatant (Figure 5C). These findings strongly suggest that the enhanced cellular LDL uptake induced by EL overexpression in LDLR deficient hepatocytes is mediated by EL’s enzymatic activity.

**Figure 4.**
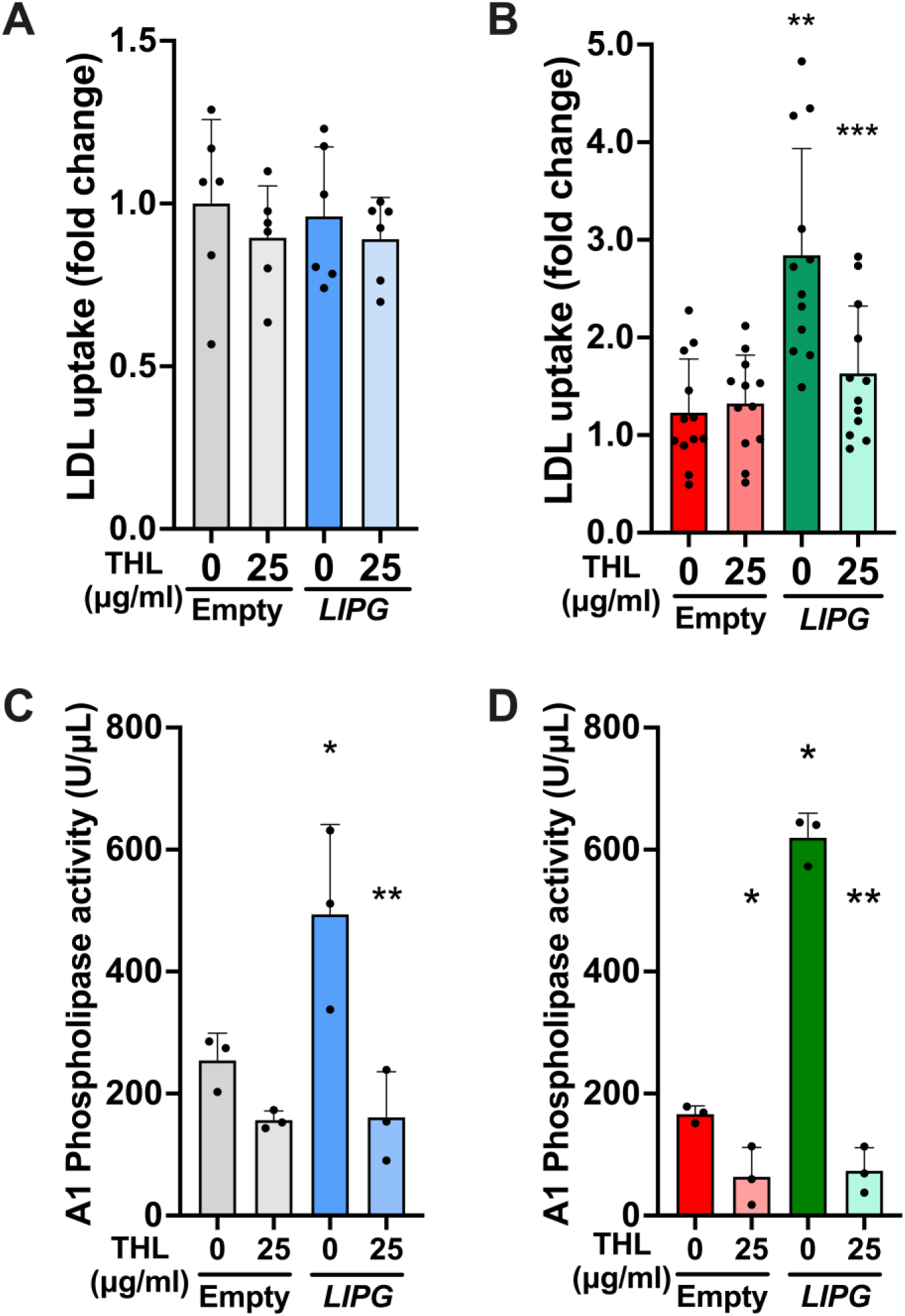
Treatment with the lipase inhibitor tetrahydrolipstatin (THL) diminished the LIPG-induced boost in cellular LDL uptake in LDLR-KO hepatoma cells. A and B: Cellular LDL uptake in Control (A) and LDLR-KO (B) HepG2 cells, with or without EL overexpression (LIPG), in the presence or absence of THL. Cells were transfected with either an empty plasmid or a plasmid containing the human LIPG gene (EL). After 72 h, cells were incubated with vehicle (DMSO) or 25 µg/ml THL (Sigma Aldrich, St. Louis, MO) for 3 hours. Then, cells were incubated with 20 µg/ml of fluorescently labeled human LDL for 90 minutes in the presence of vehicle or 25 µg/ml THL. Cellular uptake of fluorescent LDL was measured by FACS. C and D. A1 phospholipase activity in post-heparin conditioned media from control (C) and LDLR-KO (D) HepG2 cells. Cells, with and without EL overexpression, were treated with vehicle (DMSO) or 25 µg/ml THL (Sigma Aldrich, St. Louis, MO) for 3 hours at 37°C. Then, cells were incubated with 5 U/mL heparin in Opti-MEM^TM^ Serum-free media (Gibco, Waltham, MA) for 90 minutes at 37°C, in the presence of vehicle or 25 µg/ml THL. A1 phospholipase activity was measured in the conditioned media with a commercial kit (EnzChek™ Phospholipase A1 Assay Kit, Invitrogen, Waltham, MA). Data is presented as mean ± SD. One Way-ANOVA. *p<0.04 vs. cells transfected with an empty plasmid plus vehicle (0 ug/ml THL). **p<0.01 vs. LIPG-overexpressing cells plus vehicle (0 ug/ml THL).

**Figure 5.**
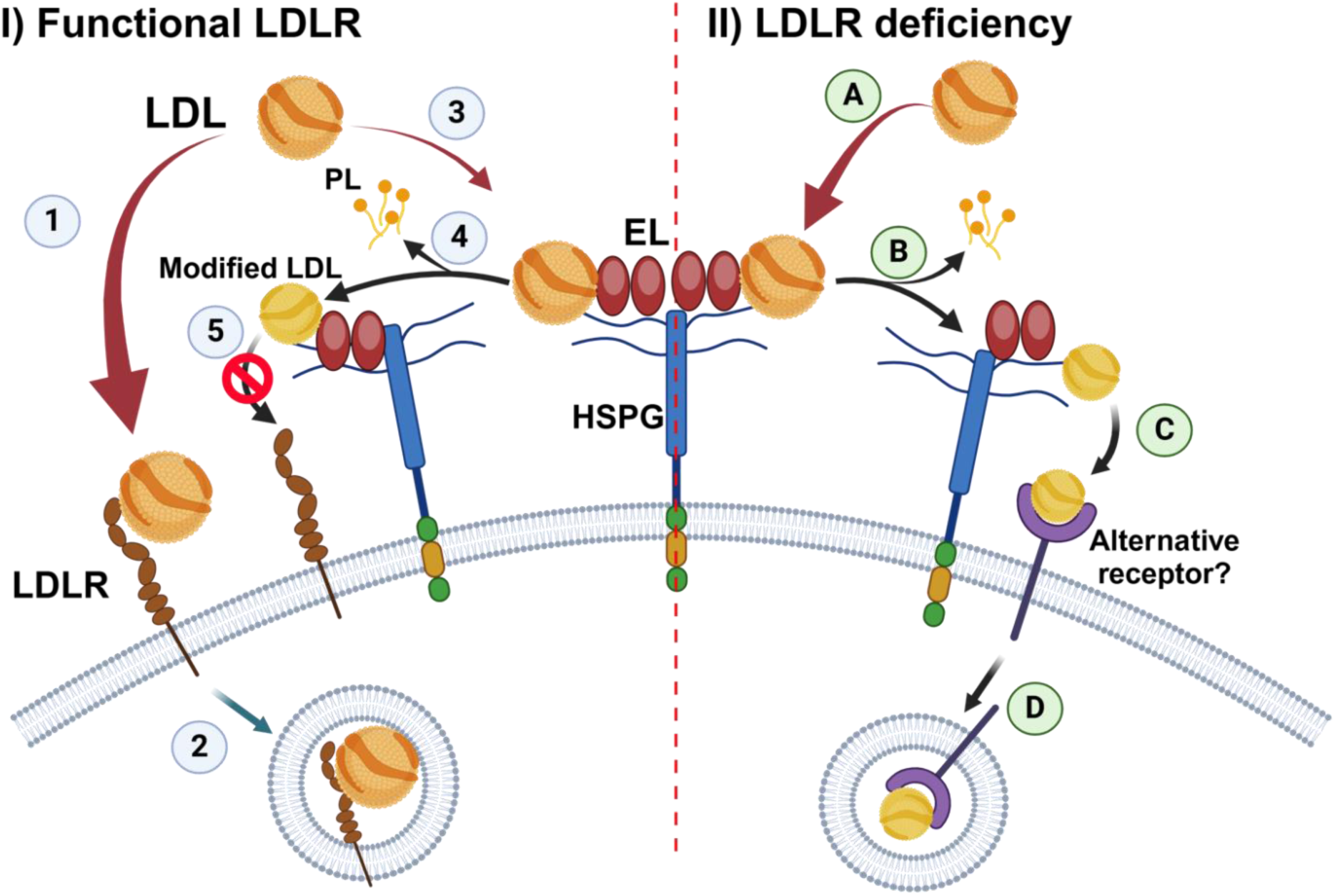
Proposed model for endothelial lipase’s role in cellular LDL uptake. *In the presence of functional LDLR (left):* 1) LDL binds with high affinity to the LDLR. 2) The LDL/LDLR complex is internalized. 3) A small portion of LDL interacts with the HSPG/EL complex. 4) EL modifies LDL by remodeling its phospholipids (PL). 5) Modified LDL is poorly recognized by the LDLR, which leads to a slight drop in LDL uptake in the control cells overexpressing EL (Fig. 2B). *In LDLR deficiency (right):* A) LDL interacts with the HSPG/EL complex. B) EL’s phospholipase activity modifies LDL composition by removing PL. C) Modified LDL becomes a better ligand for an alternative receptor that might be upregulated in LDLR deficiency. D) The LDL/alternative receptor complex is internalized into the cells.

## DISCUSSION

Although LDLR deficiency in mice and humans leads to marked increases in circulating LDL, the fact that LDL typically does not increase more than 3-4 fold in LDLR-deficient patients and mice indicates that there must be alternative pathways for LDL removal from the circulation. Other authors have reported that, in *Ldlr*-KO mice, 40% of plasma LDL is cleared from the circulation in 4 hours (27). In addition, here, we report that after knocking out the LDLR in hepatoma cells, there is a remaining 20% of residual LDL uptake, which further supports the existence of LDLR-independent mechanisms for hepatic LDL internalization. In fact, the administration of ANGPTL3 inhibitors to LDLR-null FH patients resulted in a strong reduction in LDL cholesterol (5). These findings confirm the hypothesis of a LDLR-independent pathway for LDL catabolism which can be therapeutically targeted to reduce LDL in high-risk patients. Notably, recent data suggest that EL may be involved in this pathway (12), but its role remains unclear. Here, we used LDLR-deficient hepatocytes to explore the role of EL in cellular LDL uptake. We found that overexpressing EL in LDLR-deficient hepatocytes enhanced the cellular LDL uptake through an LDLR-independent pathway that relies on EL’s enzymatic activity. This pathway, which seems irrelevant in LDLR sufficient cells, may provide an alternative explanation for the LDL-lowering effect of ANGPTL3 inhibitors in FH.

Early studies reported *LIPG* mRNA expression in the liver by northern blot analysis of whole tissue (26), but these analyses did not account for cell-specific EL expression. Here, we show that hepatocytes express *LIPG* mRNA at significant levels. In addition, our analysis of *LDLR*-KO hepatoma cells showed greater *LIPG* mRNA expression compared to controls, possibly because of compensatory mechanisms aimed to increase LDL uptake. Nonetheless, further studies will elucidate whether the upregulation of hepatocyte-specific EL expression is solely responsible for LDLR-independent LDL catabolism *in vivo*.

One of the main contributions of our study is that the effects of EL on LDL uptake are exclusive to *LDLR*-KO cells. In LDLR sufficiency, EL does not play a significant role in the LDLR-mediated uptake but rather it slightly interferes with this pathway, possibly by modifying LDL particles and reducing their affinity for the LDLR (28). In contrast, EL-overexpression nearly fully rescued the LDL uptake in LDLR deficiency by enhancing the rate of LDL internalization. Post-heparin lipases, especially lipoprotein lipase (LPL), have been shown to act as a biochemical bridge between HSPGs and lipoproteins (29–31). However, in the case of EL, our experiments with THL demonstrate that EL enzymatic activity is required for the EL-mediated, LDLR-independent LDL uptake in hepatocytes without changes in the overall LDL binding. Our studies using heparin and heparinases suggest that the EL enzymatic actions must occur in proximity to the hepatocyte surface or that the uptake pathway requires cell surface HSPGs. These findings align with previous observations that the overexpression of only enzymatically active EL accelerates LDL catabolism in LDLR deficient mice (17). Nevertheless, others have reported increased LDL binding and uptake in CHO cells overexpressing catalytically inactive EL (32). Discrepancies with our findings may be due to the use of different cell types; however, we cannot rule out the possibility that EL overexpression enhances LDL uptake via a combination of enzymatic and non-enzymatic mechanisms.

It is possible that, in LDLR deficiency, EL-mediated modifications on LDL may favor LDL uptake through alternative pathways. In fact, EL overexpression *in vivo* promotes the formation of lipid-depleted, smaller LDL particles (17), which bind to HSPG with greater affinity than mid-size LDL particles (33). Here, two different conventional methods to modulate HSPG metabolism, namely the addition of low dose heparin and heparinases, revealed that HSPGs play a role not only in EL-mediated LDL uptake but also in LDLR-independent LDL uptake in general. Interestingly, the deletion of syndecan 1, which mediates chylomicron clearance (34), does not affect the reduction of LDL induced by ANGPTL3 inhibitors (12). Therefore, the specific HSPG that mediates LDLR-independent LDL uptake remains to be determined.

It is generally believed that hepatic HSPGs represent a low-affinity, high-capacity lipoprotein binding system (34, 35). However, the saturation of EL-mediated LDL uptake at modest LDL concentrations in *LDLR*-KO cells, reported here, contrasts with this model. These results indicate the possible participation of an alternative receptor that may bind EL-modified LDL particles but would saturate at excess of LDL. This is further supported by the inhibition of the uptake due to an excess of unlabeled LDL, like LDLR-mediated uptake. Possible alternative receptors include scavenger receptors with high affinity for modified LDL particles. Interestingly, others have deleted the expression of scavenger receptor class B type I (SR-BI) in LDLR deficient mice and shown that this receptor is not involved in the reduction of LDL induced by ANGPTL3 inhibitors (12). Therefore, further studies will be needed to elucidate which alternative receptors are involved in the EL-mediated LDL uptake.

This study has several potential limitations. The *in vitro* design of our study limits the extrapolation of our conclusions to FH patients; however, the mechanisms proposed here are in line with data obtained in animal studies (12, 17) and the clinical observations in FH patients treated with ANGPTL3 inhibitors (5). Furthermore, we used THL, a widely used panlipase inhibitor, to test the influence of EL activity on cellular LDL uptake. No cytotoxicity was observed with the chosen concentration (data not shown); however, we cannot exclude potential unaccounted side effects on cells such as changes in the rate of endocytosis. Future studies will implement catalytically inactive EL. In addition, we did not assess EL’s effects on LDL composition. Future studies will address how the specific remodeling induced by EL affects LDL cellular uptake. Finally, ANGPTL3 inhibitors were not used in our studies. It should be noted that ANGPTL3 deletion appears to influence multiple aspects of lipid metabolism (19), which could potentially confound the data interpretation. By focusing on EL’s effects, our study explored EL’s impact on the hepatic uptake of LDL, a key step of LDL metabolism, without the pleiotropic influence of ANGPTL3 inhibition. Even so, our findings contribute to the general understanding of the complex mechanisms that mediate lipid-lowering effects of ANGPTL3 inhibitors.

In summary, we demonstrate that EL overexpression enhances the internalization of LDL in LDLR-deficient hepatoma cells through a mechanism that requires EL’s enzymatic activity. Our data are consistent with studies by others suggesting that EL’s phospholipase activity changes LDL composition, making the modified LDL a better ligand for alternative internalization mechanisms, namely HSPGs or unidentified alternative receptors (Figure 5). Importantly, these mechanisms are seemingly upregulated in LDLR deficiency and inactive in LDLR sufficiency. Here, we present a mechanism that, in synergism with the previously proposed (12), could explain the kinetics of the EL-dependent reduction of LDL cholesterol observed in FH patients treated with ANGPTL3 inhibitors (18). The most widely used strategies to lower plasma LDL, such as the use of statins and PCSK9 inhibitors, require the upregulation of LDLR. The pathways illustrated in our study may play a key role in lowering LDL of patients with LDLR deficiency; however, a better understanding of these alternative pathways could allow for the development of new lipid-lowering therapies that may also be used in the general population.

## ACKNOWLEDGEMENTS

We would also like to thank to the Flow Cytometry, NHLBI, at the National Institutes of Health, for their technical support.

## FUNDING SOURCES

This research was supported by the Intramural Research Program of the National Heart, Lung, and Blood Institute (NHLBI) (HL006095) at National Institutes of Health and the Leducq Foundation (Grant: 23CVD02) (AR). I.J.G. and N.O.S. are supported in part by grant P01 HL151328 from the National Heart, Lung, and Blood Institute (NHLBI).

## Notes

### Competing Interest Statement

The authors have declared no competing interest.

### Summary of Updates

The focus of the introduction and discussion was tightened on endothelial lipase.

